# Cortical Iron in Schizophrenia: A Quantitative Susceptibility Mapping and Diffusion Tensor Imaging MRI Study

**DOI:** 10.1101/2025.10.17.683066

**Authors:** Luke James Vano, Jan Sedlacik, Stephen John Kaar, Grazia Rutigliano, Richard Carr, Alaine Berry, Ben Statton, Amir Fazlollahi, Oliver David Howes, Robert Ali McCutcheon

## Abstract

**Background and Hypothesis:** Cognitive and negative symptoms in schizophrenia remain poorly treated. Iron dysregulation has been implicated as a potential mechanism underlying cognitive dysfunction and schizophrenia. While elevated postmortem iron in Brodmann areas 10-11 has been linked to schizophrenia, this has not been assessed *in vivo*. We therefore used iron-sensitive MRI to test whether cortical iron is elevated in individuals with schizophrenia compared to healthy controls.

**Study Design:** We acquired quantitative susceptibility mapping (QSM) MRI to measure magnetic susceptibility (χ), a marker of iron, in 158 participants aged 18–45 (73 with schizophrenia and 76 matched healthy controls). As χ is reduced by myelin, we conducted diffusion tensor imaging (DTI) to assess mean diffusivity, an iron-insensitive marker also reduced by myelin.

**Study Results:** Primary analyses showed no significant case-control differences in χ in the whole cortex (p=0.675) or Brodmann areas 10–11 (p=0.537). Exploratory analyses examined χ for 362 cortical regions and a voxelwise analysis, correcting for multiple comparisons. Two left temporo-parieto-occipital (TPO) junction regions showed significantly elevated χ in schizophrenia: the posterior TPO junction (d=0.752, p<0.001) and the superior temporal visual area (d=0.638, p=0.033), which remained significant after adjusting for mean diffusivity and clinical covariates (p=0.001 and p=0.023, respectively). Voxelwise analysis confirmed elevated χ in schizophrenia in the left TPO junction (peak t=5.62).

**Conclusions:** This study provides the first *in vivo* evidence of elevated cortical iron in schizophrenia, suggesting regional iron accumulation may contribute to cortical pathology.

## Introduction

While antipsychotic medications can treat positive psychotic symptoms of schizophrenia, they are minimally effective at improving cognitive (1) or negative symptoms (2). Since these deficits have been linked to cortical dysfunction (3), identifying the underlying biology of disease-related cortical abnormalities could inform the development of new therapeutic strategies.

Iron is essential for many neurophysiological processes including myelination, neurotransmitter synthesis, and neurogenesis (1). Iron deficiency is one of the leading causes of impaired cognitive development globally (4). Iron levels must be tightly regulated, as excess brain iron can damage grey matter through increased oxidative stress, lipid peroxidation, and neuroinflammation (5). A recent postmortem study of Brodmann areas 10-11 reported elevated iron but reduced ferritin—the major storage protein of iron—in schizophrenia (6). Because ferritin normally sequesters iron in a redox-inactive form, it was hypothesised that this may indicate an increased pool of chemically active iron, leading to the cortical thinning observed on structural MRI in schizophrenia (7,8).

Quantitative susceptibility mapping (QSM) is an MRI technique that estimates tissue magnetic susceptibility (χ), which is increased by iron and decreased by myelin (9). Previous case-control studies have linked early-course schizophrenia to lower subcortical χ (10–13), suggesting disease-related iron loss or myelin gain. Recently, using diffusion tensor imaging (DTI) alongside QSM, we demonstrated that this χ reduction was associated with elevated subcortical mean diffusivity (13), an iron-insensitive marker that is reduced by myelin (14).

These findings indicate that lower iron, rather than higher myelin, underlies the lower subcortical χ observed in schizophrenia. As *in vivo* measurement of cortical iron in schizophrenia has not previously been reported, we extended our investigation to examine cortical QSM and DTI in the same cohort.

## Materials and Methods

### Participants

Ethical approval was granted by the London-Dulwich Research Ethics Committee granted ethical approval for healthy participants and by the Office for Research Ethics Committees Northern Ireland for individuals with schizophrenia (NCT04038957). These participants also underwent neuromelanin-sensitive MRI and [18F]-DOPA positron emission tomography as part of previously published studies (12,15). All participants provided written informed consent. Recruitment was limited to individuals aged 18–45 years, to target those in the early stages of illness.

Individuals diagnosed with schizophrenia were referred from community mental health services in London, UK. Diagnoses were confirmed by a study psychiatrist using clinical records and the Structured Clinical Interview for DSM-5 (SCID-5) (16). Participants were excluded if they had comorbid psychiatric diagnoses, current or past substance use disorders (excluding nicotine), were taking non-antipsychotic psychotropic medications, more than one antipsychotic, or current or prior clozapine treatment.

Healthy control participants were recruited through public advertisements in London, UK. A psychiatrist screened volunteers, excluding those with personal psychiatric history, psychotropic use, substance use disorder (excluding nicotine), or a first-degree relative with schizophrenia.

All participants underwent a urine drug screen (UDS) prior to their MRI. Individuals testing positive for stimulants, benzodiazepines, or opiates were excluded. Participants who were positive for delta-9-tetrahydrocannabinol (THC), which may persist in urine for extended periods (17), were able to participate as long as they had not used it in the past month. Smoking status was recorded as current, former, or never.

### Clinical Assessments

Demographic and clinical data were collected from all participants. Patients completed the Positive and Negative Syndrome Scale (PANSS) (18), the Brief Negative Symptom Scale (BNSS) (19), and the Clinical Global Impression-Severity scale (CGI-S) (20). Patient antipsychotic doses were converted to chlorpromazine daily equivalents (21).

### MRI Acquisition

We scanned participants using a 3T MRI scanner (Magnetom Prisma, Siemens Healthineers, Erlangen, Germany) with a 64-channel receive head coil. Longitudinal relaxation time weighted (T1w) images were generated with a magnetization prepared rapid gradient echo (MPRAGE; acquisition time (TA)=5 mins 35 secs, repetition time (TR)=2300ms, inversion time (TI)=900ms, echo time (TE)=2.91ms, flip angle=9°, 176 slices with the voxel size=1×1×1mm^3^).

QSM maps were calculated from magnitude and phase images generated from a 3D gradient recall echo (GRE) sequence (TA=8 mins 5 secs, TR=50ms, first TE at 5.84ms with 8 subsequent echoes 4.79ms apart, flip angle=15°, 144 slices with the voxel size=1×1x1mm^3^). QSM images were quality controlled by investigators LJV and JS, who were blinded to diagnosis and used best practice guidelines (22). χ is reported in parts per billion (ppb).

DTI was acquired using a single-shot echo-planar imaging sequence (TA=7 mins 36 secs; TR=3200 ms; TE=69 ms), comprising 27 axial slices with a voxel size of 1.7×1.7×4 mm^3^. Eleven non-diffusion-weighted (b=0 s/mm^2^) images and 2 diffusion-weighted images (b=1000 s/mm^2^) with 64 uniformly distributed gradient directions were collected. An additional sequence with reversed phase-encoding direction and no diffusion weighting was acquired using otherwise identical parameters for distortion correction during preprocessing.

### QSM Preprocessing

QSM maps can be distorted near B0 inhomogeneities, such as air-tissue or bone-tissue boundaries next to the orbitalfrontal cortex and inferior temporal lobes. These areas exhibit signal drop-out on GRE magnitude images and are typically excluded by conservative brain masking. Therefore, we generated a brain mask from the first-echo magnitude image using the FMRIB Software Laboratory (FSL) Brain Extraction Tool (BET), which we then eroded by 5% (23). As this mask lacks precision at cortical boundaries, FreeSurfer was used to generate a Desikan-Killiany-Tourville cortical atlas on T1w images (24). The GRE magnitude image was co-registered to the T1w image using NiftyReg with a 15 mm spline grid to restrict non-linear warping to regions with substantial distortion (25). The resulting transformations were used to map the FreeSurfer mask into GRE space. QSM processing was limited to voxels within both masks.

Frequency shift was computed from brain-masked phase images acquired with the GRE sequence, using Fit_ppm_complex.m from the Morphology Enabled Dipole Inversion (MEDI) toolbox (26). Local frequency was estimated via projection onto dipole fields (27), and QSM maps were generated using iterative Tikhonov dipole inversion (28). Only cortical voxels with Desikan-Killiany-Tourville labels >1000, as well as hippocampal (17, 53) and amygdala (18, 54) voxels, were retained.

For our region-of-interest (ROI) analysis, we employed the extended Human Connectome Project parcellation (HCPex v1.1) (29). As this atlas is in the Montreal Neuroimaging Institute (MNI) space, we used sMRIprep (v0.17.0) to normalise the T1w images (30). The relevant inverse affine transformation-matrices and warp-fields were applied to move the atlas from the MNI to GRE space. Amygdala labels (387, 420) were added to the 360 cortical HCPex labels. FreeSurfer cortical voxels were re-labelled based on overlapping HCPex atlas voxels or nearest-neighbour assignment. ROIs corresponding to Brodmann areas 10-11 (labels 134, 135, 150, 151, 156, 158, 314, 315, 316, 330, 336, 338, 148, 152, 157, 328, 332, and 337) were combined into a single mask for our primary analysis.

The size of each ROI was calculated by counting the number of voxels with each label in the cortical parcellation. We then applied the mask generated from the first-echo magnitude image, which removes voxels experiencing significant distortion, to generate our final cortical parcellated atlas. The size of each ROI in this mask was divided by the value from the original cortical parcellation to estimate the effect that distortion had on an ROI.

For voxelwise analysis, cortical QSM maps were transformed into MNI space using the relevant affine and nonlinear transformations. To minimize the influence of outliers, voxels with χ values below the 1st percentile or above the 99th percentile were excluded. Voxelwise analysis was restricted to the remaining cortical voxels, which were smoothed with a Gaussian kernel (σ=3 mm).

### DTI Preprocessing

We used FSL’s diffusion toolbox (31) to preprocess the DTI data. Images were corrected for the effects of eddy currents, distortion, and head movement (32). Following brain extraction, we used dtifit to fit a diffusion tensor model with FSL’s tract-based spatial statistics (TBSS) toolbox (33). Participant B0 data were linearly coregistered to their T1w images. Any HCPex labels or clusters showing significant case-control χ differences were moved from MNI space to T1w space and then into DTI space using the relevant inverse affine transformation-matrices and warp-fields. Mean diffusivity values within these masks were then calculated.

## Statistical Analysis

### Baseline Clinicodemographic Analysis

We examined for case-control clinicodemographic differences with chi-squared tests for categorical variables and independent sample t-tests for continuous variables.

### Case-Control QSM Analysis

Participants were included in the ROI analysis if ≥70% of voxels within the region were retained after masking. This threshold was chosen for consistency with previous cortical analyses using our earlier pipeline (34). Results were only reported if at least 50 participants remained in each group and the group difference in masked voxel count was no more than 2% for the entire ROI. For our primary analyses, we used independent sample t-tests to assess the significance and Cohen’s d for the effect size of any case-control differences in χ for the whole cortex and Brodmann 10–11 mask. Exploratory analyses examined all ROIs, with significance assessed via false discovery rate (FDR) correction using the Benjamini-Hochberg method (p<0.05). For our primary analyses, and if χ case-control differences were identified for any mask, a robust linear regression model was used to determine whether these were influenced by clinical confounders (age, sex, current or past smoking history, and THC-positive UDS) or mean diffusivity.

To identify voxels showing significant case-control differences, we applied threshold-free cluster enhancement (TFCE) with family-wise error (FWE) correction using 10,000 permutations (p<0.05) (35). The case-control differences in mean diffusivity for any clusters showing significant case-control χ differences were examined using t-tests.

## Results

### Baseline Clinicodemographic Analysis

QSM maps were generated for 171 participants (86 healthy controls and 85 with early course schizophrenia). Twenty-two of these maps failed quality control: 7 for excessive movement (3 healthy controls and 4 with schizophrenia) and 15 for artifacts (7 healthy controls and 8 with schizophrenia). The final dataset included 76 healthy controls and 73 with schizophrenia (17 were antipsychotic-free). DTI was performed in 101 participants (38 healthy controls and 63 with schizophrenia) with usable QSM data. Clinicodemographic information is reported in Table 1. There were no significant case-control differences in age, sex, ethnicity, or THC-positive status but current smoking was more prevalent in the schizophrenia group compared to controls (34% vs. 16%; p=0.03).

**Table 1.**
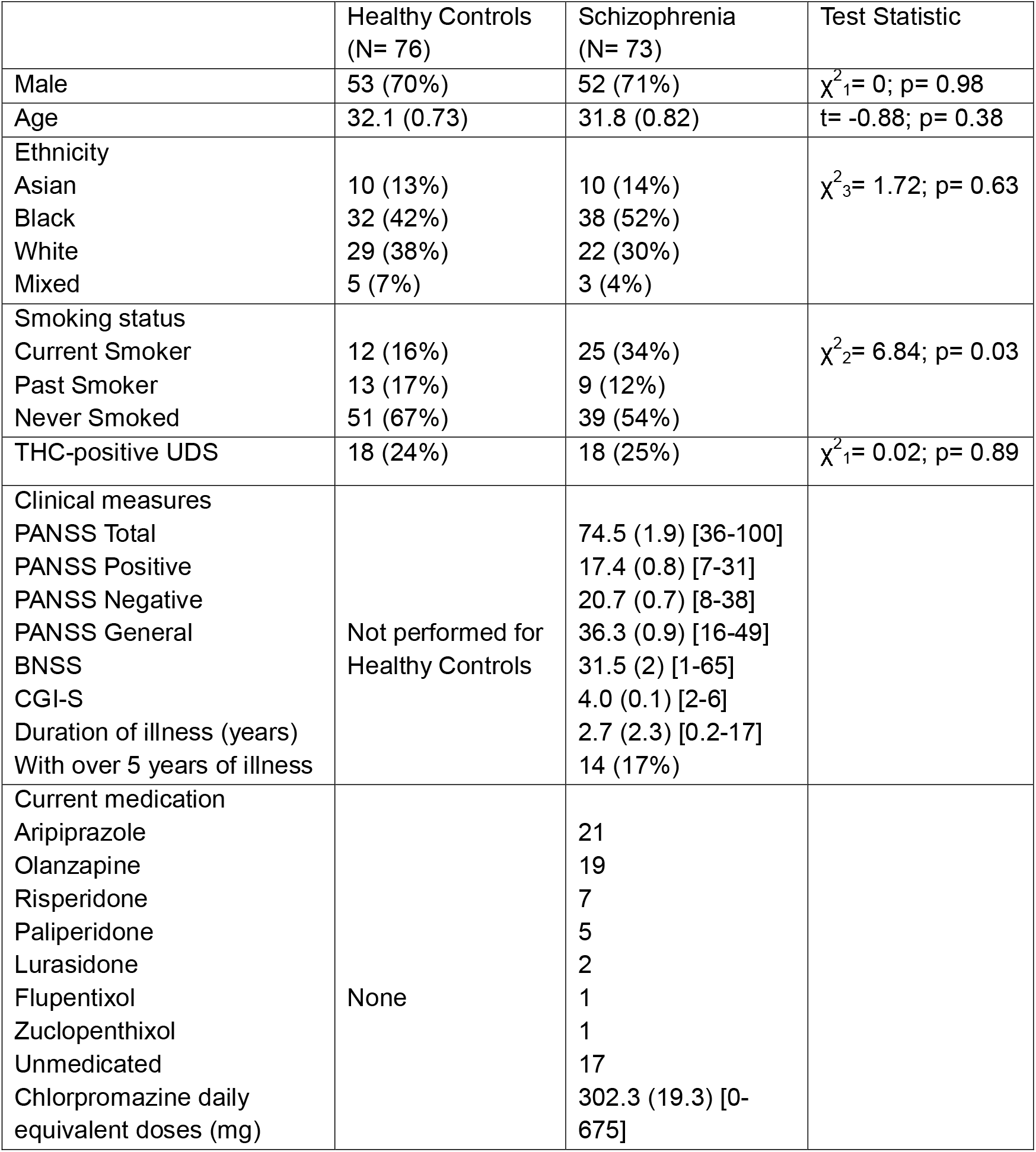
Participant clinicodemographic data of those with usable quantitative susceptibility mapping data. Values are expressed as number (and %) or mean (with standard error in brackets) [and range in square brackets], besides duration of illness that was expressed as median (and interquartile range). N, number of participants; THC-positive UDS, delta-9-tetrahydrocannabinol positive urinary drug screen; PANSS, Positive and Negative Syndrome Scale; BNSS, Brief Negative Symptoms Scale; CGI-S, Clinical Global Impression-Severity Scale; χ, chi-squared test; t, independent sample t-test.

### Case-Control QSM Analysis

Figure 1 displays the results of the case–control analyses. There was no significant difference in whole cortex χ between healthy controls (mean = 1.49, SE = 0.21) and patients with schizophrenia (mean = 1.62, SE = 0.23; d=0.07, 95% CI: [-0.25, 0.39], p=0.675). This remained non-significant in our robust regression that controlled for potential confounders and mean diffusivity (p=0.899; Table S1). Similarly, χ within the Brodmann areas 10-11 mask was not significantly different between groups in our primary analysis (mean = –2.44, SE = 0.25; mean = –2.23, SE = 0.25; d=0.07, 95% CI: [-0.25, 0.39], p=0.537) or when controlling for potential confounders and mean diffusivity (p=0.763; Table S2).

**Figure 1.**
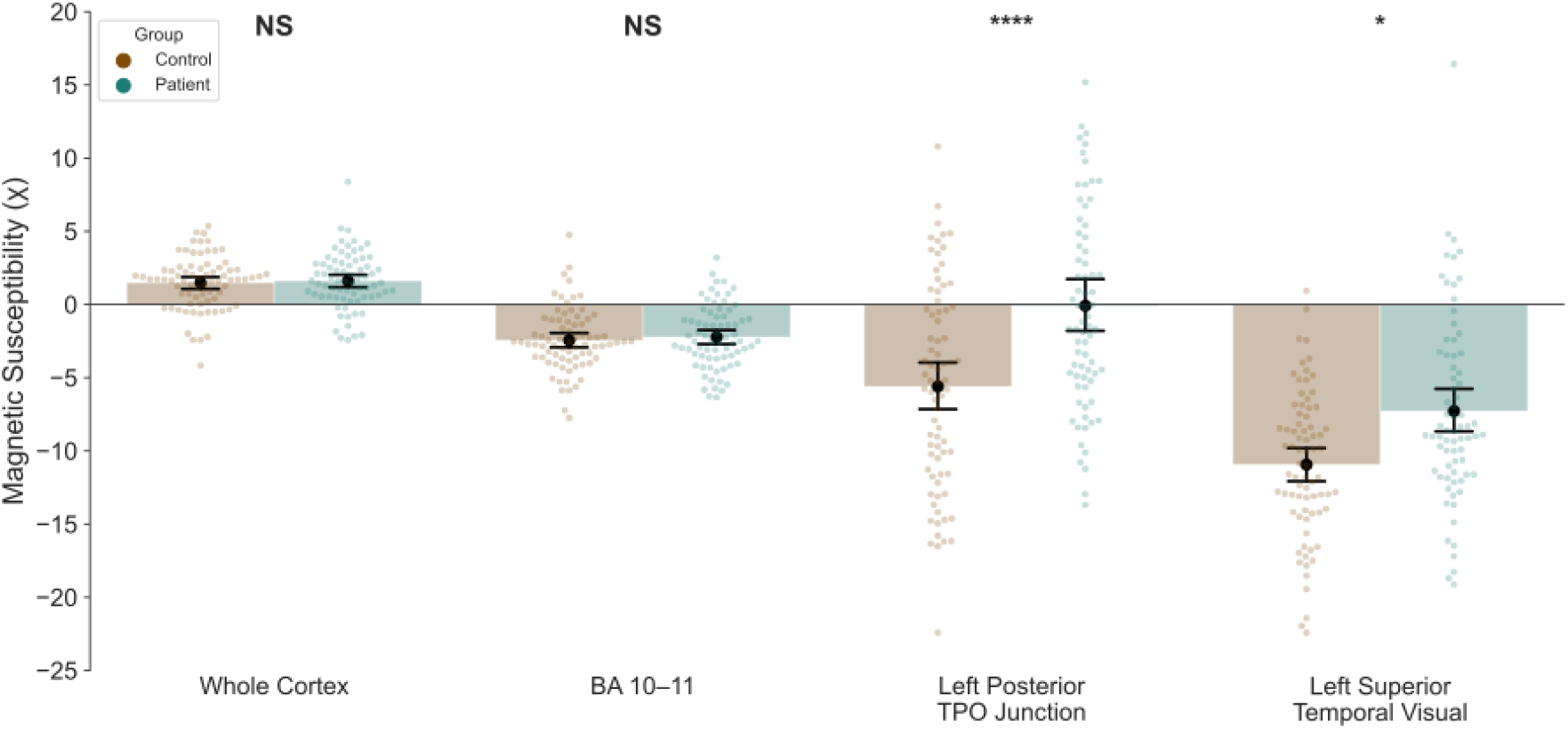
Group differences in magnetic susceptibility (χ) across selected cortical regions. Bars show group means with standard error, overlaid with individual participant values. Data from healthy controls are shown in brown; data from patients with schizophrenia are shown in green. In the predefined regions of interest, there were no significant differences in χ between groups for the whole cortex (p = 0.675) or Brodmann areas (BA) 10–11 (p = 0.537). In exploratory analyses, χ was significantly elevated in schizophrenia after false discovery rate correction in two left temporo-parieto-occipital (TPO) junction regions: the posterior third of the TPO junction (p < 0.0001) and the left superior temporal visual area (p = 0.033).

Case-control results for the cortical ROIs are presented in Supplementary File 1. Two ROIs within the left temporo-parieto-occipital (TPO) junction remained significantly different after FDR correction, both showing elevated χ in schizophrenia. These were the TPO junction area 3, which comprises the posterior part of the TPO junction (d=0.752, 95% CI: [0.419, 1.084], FDR-corrected p<0.0001), and the superior temporal visual area (d=0.638, 95% CI: [0.308, 0.967], FDR-corrected p=0.033). These findings remained significant when controlling for potential confounders and mean diffusivity in the TPO junction area 3 (z=3.19, p=0.001; Table 2) and superior temporal visual area (z=2.27, p=0.023; Table 3).

**Table 2.** Results from the robust linear model built to predict left temporo-parieto-occipital (TPO) junction area 3 magnetic susceptibility (χ) with case-control status, potential confounders, and mean diffusivity.

| Variable | Coefficient | Standard Error | t-score | p-value | [95% CI] |  |
| --- | --- | --- | --- | --- | --- | --- |
| (Intercept) | 10.16 | 13.44 | 0.76 | 0.45 | -16.18 | 36.50 |
| Schizophrenia Group Status | 4.90 | 1.54 | 3.19 | 0.001 | 1.89 | 7.92 |
| Current Smoker | -1.16 | 1.81 | -0.64 | 0.522 | -4.71 | 2.39 |
| Past Smoker | 2.92 | 2.09 | 1.40 | 0.162 | -1.17 | 7.01 |
| THC-positive UDS | 0.31 | 1.76 | 0.18 | 0.858 | -3.13 | 3.76 |
| Male Sex | -1.18 | 1.65 | -0.72 | 0.474 | -4.41 | 2.05 |
| Age | 0.05 | 0.11 | 0.50 | 0.616 | -0.16 | 0.26 |
| Mean Diffusivity | -17.89 | 14.59 | -1.23 | 0.22 | -46.49 | 10.72 |

**Table 3.** Results from the robust linear model built to predict left superior temporal visual area magnetic susceptibility (χ) with case-control status, potential confounders, and mean diffusivity.

| Variable | Coefficient | Standard Error | t-score | p-value | [95% CI] |  |
| --- | --- | --- | --- | --- | --- | --- |
| (Intercept) | -21.06 | 13.27 | -1.59 | 0.113 | -47.06 | 4.95 |
| Schizophrenia Group Status | 2.84 | 1.25 | 2.27 | 0.023 | 0.39 | 5.30 |
| Current Smoker | 2.95 | 1.48 | 2.00 | 0.046 | 0.06 | 5.85 |
| Past Smoker | 2.06 | 1.69 | 1.22 | 0.221 | -1.24 | 5.37 |
| THC-positive UDS | -1.20 | 1.44 | -0.84 | 0.403 | -4.03 | 1.62 |
| Male Sex | -0.48 | 1.35 | -0.36 | 0.721 | -3.12 | 2.16 |
| Age | 0.05 | 0.09 | 0.63 | 0.53 | -0.11 | 0.22 |
| Mean Diffusivity | 8.78 | 14.29 | 0.62 | 0.539 | -19.23 | 36.79 |

Our voxelwise analysis identified a cluster within this region where schizophrenia was associated with greater χ (size=1261 voxels, peak t-value = 5.62, FWE p<0.05; Figure 2). Mean diffusivity within this cluster was not significantly different between groups (t=0.42, p=0.676). There were no significant clusters where χ was lower in schizophrenia.

**Figure 2.**
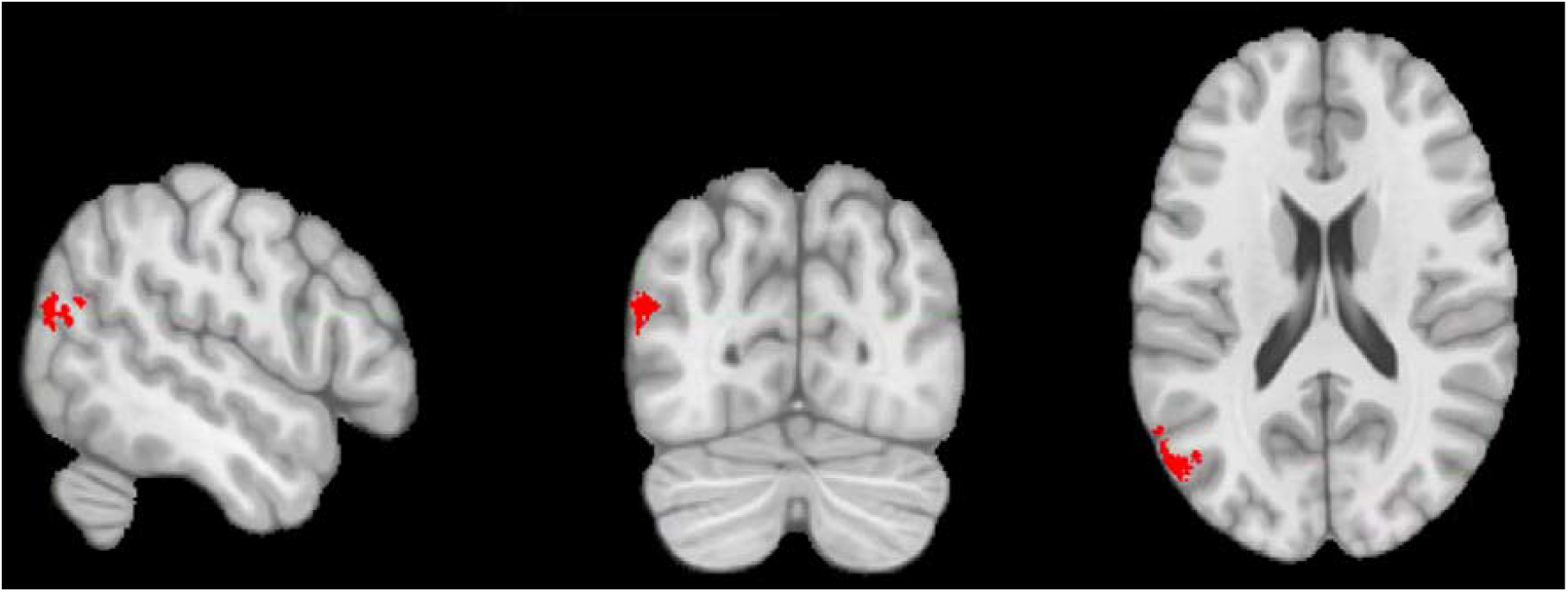
Voxelwise cluster within the left temporo-parieto-occipital junction showing significantly greater magnetic susceptibility (χ) in patients with schizophrenia compared to matched healthy controls. The cluster (shown in red) comprised 1261 voxels, with the peak voxel (t-value = 5.62) at Montreal Neuroimaging Institute coordinates x = –54, y = –73, z = 20. This result survived family-wise error correction (p < 0.05).

## Discussion

Using QSM we present, to our knowledge, the first *in vivo* investigation of case-control cortical iron differences in schizophrenia. While prior postmortem work reported elevated iron in Brodmann areas 10-11 in schizophrenia (6), we found no case-control differences in χ in these regions or when examining the whole cortex. However, our exploratory analyses identified two ROIs, and one voxel cluster within the left TPO junction with significantly higher χ in schizophrenia. As DTI-derived mean diffusivity, which is lowered by myelin, did not mediate the χ group difference, this is more likely explained by higher iron in schizophrenia.

Notably, these results build upon previous findings from the same participant cohort, where patients with schizophrenia showed higher neuromelanin-sensitive MRI signal (15)—a marker of neuromelanin-bound iron—and lower χ in the substantia nigra and ventral tegmental area (12). Both factors were independently linked to higher striatal dopamine synthesis capacity in the patient group. Moreover, subcortical χ was reduced and mean diffusivity elevated in schizophrenia (13), supporting the interpretation that reduced iron, rather than increased myelin, underlies subcortical χ differences. Collectively, our findings suggest that schizophrenia is characterized by regionally specific disruptions in brain iron homeostasis.

### Methodological Considerations

Accurately measuring cortical χ with QSM is technically challenging, particularly in areas prone to distortion, such as the orbitofrontal and inferior temporal cortices (22). We minimized the impact of distortion-related susceptibility artifacts by excluding voxels with extreme values and only including ROIs where a majority of voxels passed quality control. Dual masking with both GRE magnitude and T1w images increased our ability to remove artifact-prone areas. However, residual susceptibility artifacts likely reduced sensitivity in some cortical areas—particularly the orbitofrontal cortex—which may explain the absence of a case-control difference in Brodmann areas 10–11, despite postmortem evidence of increased iron here in schizophrenia (6).

Previous studies have identified raised T1w/T2w ratios in the left TPO junction in schizophrenia compared to controls, despite lower ratios in other cortical ROIs (36,37). Although initially proposed as a proxy for intracortical myelin, postmortem validation shows that T1w/T2w ratios negatively correlate with myelin and positively with χ, suggesting that they are iron-sensitive (38). This is supported by the highest T1w/T2w ratios occurring in iron-rich, relatively myelin-poor subcortical regions (38,39). Thus, we interpret that higher χ and T1w/T2w ratios within the left TPO junction reflect iron accumulation. Elevated myelin is unlikely to explain our findings, given that mean diffusivity (which inversely correlates with myelin) did not mediate higher left TPO junction χ in schizophrenia. Additionally, we previously showed higher subcortical mean diffusivity and lower white matter magnetic susceptibility anisotropy (an iron-insensitive myelin marker) in this schizophrenia cohort relative to controls (13), suggesting that brain myelin is reduced in our participants with schizophrenia.

### Implications for Understanding Iron Homeostasis in Schizophrenia

Although peripheral iron deficiency is associated with schizophrenia, central and peripheral iron homeostasis are largely independent due to the blood-brain barrier (5). Findings from previous subcortical case-control iron-sensitive MRI studies have supported both high (40,41) and low (10,11,42) iron in schizophrenia. Our prior subcortical analysis of this participant cohort showed that subcortical χ was lower in schizophrenia (13). Therefore, our current finding that χ was increased in participants with schizophrenia within the left TPO junction but not in other cortical ROIs suggests that disease-related iron changes are regionally specific.

Elevated iron can drive neuroinflammation, oxidative stress, and lipid peroxidation, contributing to neurodegeneration (5). Cortical thinning (7,8) and progressive grey matter loss (43,44) are well-documented in schizophrenia, and our results raise the possibility that focal iron accumulation may contribute to this pathology as expertly discussed by Lotan et al (6).

The TPO junction—which lies at the intersection of the temporal, parietal, and occipital lobes—is a hub for multisensory integration (45). Voxelwise analyses with structural MRI have linked grey matter loss in this region to schizophrenia (46), particularly early-onset illness (47). Functional imaging studies report both hyperactivation and hypoactivation in the left temporoparietal subregion, with altered connectivity correlating with the severity of auditory hallucinations (48–50). Neuromodulation targeting this region, such as transcranial direct current stimulation, has shown benefits for negative symptoms and hallucinations in schizophrenia (50,50). Our finding of disrupted iron homeostasis in this area suggests a potential link between neurochemical and structural alterations, warranting further investigation.

### Limitations and future directions

Although we applied FDR correction in our exploratory analysis, the risk of type I error remains. Replication in independent samples is needed to confirm whether χ is elevated in the left TPO junction in schizophrenia. Furthermore, the application of FDR correction across 331 ROIs that passed quality control indicates a high risk of type II error. Performing this analysis on a larger dataset is necessary to examine whether case-control differences of a lower effect size exist for other ROIs.

### Conclusion

In summary, we found localized increases in iron-sensitive QSM MRI signal within the left TPO junction in schizophrenia, which were independent of group differences in myelin-sensitive mean diffusivity. This suggests that regional iron accumulation may contribute to cortical pathology in this disorder. These findings provide a foundation for further research into cortical iron as a potential biomarker and therapeutic target in schizophrenia.

## Acknowledgements

For the purpose of open access, this paper has been published under a creative common license (CC-BY) to any accepted author manuscript version arising from this submission. We thank all participants who volunteered their time to take part in the study and the community mental health teams who discussed our study with service users.

## Funding

This work was supported by Medical Research Council-UK (Grants Nos. MC_U120097115, MR/W005557/1, and MR/V013734/1 [to ODH]); Wellcome Trust (Grant No. 094849/Z/10/Z [to ODH]); the National Institute for Health and Care Research (NIHR) Biomedical Research Centre at South London and Maudsley NHS Foundation Trust and King’s College London and Sumitomo Pharma America, Inc. (study reference: NCT04038957). RAM’s work is funded by a Wellcome Trust Clinical Research Career Development Fellowship (224625/Z/21/Z). RAM is supported by the NIHR Oxford Health Biomedical Research Centre. The views expressed are those of the authors and not necessarily those of the NIHR or the Department of Health and Social Care.

## Competing Interests

RAM has received speaker/consultancy fees from Karuna, Janssen, Viatris, Boehringer Ingelheim, and Otsuka and co-directs a company that designs digital resources to support treatment of mental illness. GR is supported by the EU Horizon 2020 Research and Innovation Program under Marie Skłodowska-Curie Grant agreement no. 101026235 and by a Guarantors of Brain Post-Doctoral Clinical Fellowship and has been a consultant for Sumitomo Pharma. ODH has received investigator-initiated research funding from and/or participated in advisory/speaker meetings organized by Angellini, Autifony, Biogen, Boehringer Ingelheim, Eli Lilly, Elysium, Heptares, Global Medical Education, Invicro, Jansenn, Karuna, Lundbeck, Merck, Neurocrine, Ontrack/Pangea, Otsuka, Sunovion, Recordati, Roche, Rovi, and Viatris/Mylan. He was previously a part-time employee of Lundbeck A/v. ODH has a patent for the use of dopaminergic imaging. All other authors report no biomedical financial interests or potential conflicts of interest.

## Author Contributions

Conceptualization: L.J.V., R.A.M., S.J.K., and O.D.H. Data collection: L.J.V., G.R., S.J.K., A.B., and B.S. Formal analysis: L.J.V., R.A.M., J.S., A.B., B.S., R.C., and O.D.H. Funding acquisition: O.D.H. Methodology: L.J.V., R.A.M., J.S., S.J.K., A.F., and O.D.H. Software: L.J.V., R.A.M., J.S. Supervision: R.A.M., J.S., S.J.K., A.F., and O.D.H. Visualization: L.J.V. and R.A.M. Writing—original draft: all authors.

