## Supplemental Materials for "Cortical Iron in Schizophrenia: A Quantitative Susceptibility Mapping and Diffusion Tensor Imaging MRI Study"

### Supplementary Analyses

| Variable | Coefficient | Standard Error | t-score | p-value | [95% CI] | |
|---|---|---|---|---|---|---|
| (Intercept) | 10.09 | 7.29 | 1.38 | 0.166 | -4.20 | 24.38 |
| Schizophrenia Group Status | 0.05 | 0.37 | 0.13 | 0.899 | -0.67 | 0.77 |
| Current Smoker | 0.45 | 0.43 | 1.05 | 0.291 | -0.39 | 1.29 |
| Past Smoker | 0.53 | 0.50 | 1.06 | 0.289 | -0.45 | 1.50 |
| THC-positive UDS | -0.43 | 0.42 | -1.03 | 0.303 | -1.26 | 0.39 |
| Male Sex | 0.25 | 0.39 | 0.64 | 0.521 | -0.52 | 1.02 |
| Age | 0.09 | 0.03 | 3.62 | 0.000 | 0.04 | 0.14 |
| Mean Diffusivity | -13.17 | 8.24 | -1.60 | 0.110 | -29.32 | 2.98 |

**Table S1.** Results from the robust linear model built to predict whole cortex magnetic susceptibility ($\chi$), calculated by quantitative susceptibility mapping MRI, with case-control status, potential confounders, and diffusion tensor imaging-derived mean diffusivity.

| Variable | Coefficient | Standard Error | t-score | p-value | [95% CI] | |
|---|---|---|---|---|---|---|
| (Intercept) | -11.30 | 6.31 | -1.79 | 0.074 | -23.67 | 1.07 |
| Schizophrenia Group Status | 0.14 | 0.47 | 0.30 | 0.763 | -0.77 | 1.05 |
| Current Smoker | 0.51 | 0.54 | 0.94 | 0.345 | -0.55 | 1.58 |
| Past Smoker | 0.43 | 0.62 | 0.69 | 0.490 | -0.79 | 1.65 |
| THC-positive UDS | -0.40 | 0.53 | -0.75 | 0.455 | -1.44 | 0.65 |
| Male Sex | 0.33 | 0.50 | 0.67 | 0.504 | -0.64 | 1.30 |
| Age | 0.05 | 0.03 | 1.57 | 0.117 | -0.01 | 0.11 |
| Mean Diffusivity | 7.90 | 7.35 | 1.07 | 0.282 | -6.50 | 22.30 |

**Table S2.** Results from the robust linear model built to predict Brodmann areas (BA) 10–11 magnetic susceptibility ($\chi$), calculated by quantitative susceptibility mapping MRI, with case-control status, potential confounders, and diffusion tensor imaging-derived mean diffusivity.
